# Impaired non-shivering thermogenesis in the desert-dwelling antelope ground squirrel

**DOI:** 10.64898/2026.03.04.707825

**Authors:** Luke Olsen, Michele Albertini, Douglas Barrows, Thomas S. Carroll, Michael Hiller, Roberto Refinetti, G.J. Kenagy, Paul Cohen

## Abstract

Adaptations in animals occupying environments of extreme cold or heat offer unique insights into thermoregulation. The antelope ground squirrel, a desert-dwelling rodent situated within a clade of hibernators, provides a notable example of these thermoregulatory extremes. As a non-hibernator closely related to hibernators, antelope ground squirrels may represent a rare case of trait reversal, with a striking ability to maintain core body temperatures exceeding 43°C while displaying poor tolerance to prolonged cold. Here, we explore the genetic and phenotypic basis of these unique traits by generating the first genome assembly for this species, which we use to conduct comparative genomics across the squirrel family. We complement this with acute and chronic cold-exposure experiments coupled with transcriptomic profiling of thermogenic organs: brown adipose tissue, white adipose tissue, and skeletal muscle. Together, these findings reveal a shift away from non-shivering thermogenesis toward metabolically demanding shivering thermogenesis, a perilous strategy for sustained heat generation.

## Introduction

The dynamic control of body temperature is a critical adaptation that enables mammals to survive across the full range of earth’s climates. Small-bodied hibernators exemplify thermoregulatory extremes by adjusting their body temperature from below zero (−2.9°C) to euthermia (37°C) within hours^1^. Within the squirrel family (Sciuridae), the tribe Marmotini contains 13 genera, most of which can dynamically regulate body temperature through hibernation or estivation^2^. A notable exception is the desert-dwelling genus *Ammospermophilus* (antelope ground squirrels), which remain active year-round and do not enter these hypometabolic states^3^. Instead, antelope ground squirrels demonstrate impressive heat tolerance, with an ability to sustain core body temperatures near 43°C^4^. Intriguingly, these thermoregulatory patterns appear to have come at a cost: chronic cold exposure can be lethal, reflecting cold intolerance^5^. To date, the genomic and phenotypic signatures underpinning this unique thermal physiology remain unknown.

## Results and Discussion

To unravel the genetic determinants of thermal physiology in antelope ground squirrels, we generated the first genome for the genus *Ammospermophilus* (*A. leucurus*; Figure 1A) and the only genome for a non-hibernating member of the tribe Marmotini using long-read PacBio HiFi sequencing. The genome assembly exhibited high completeness (BUSCO Eukaryota: 94.1% complete) and strong contiguity (contig N50 = 31.58 Mb), placing it among the most contiguous marmotine genomes to date (Figure 1A and Data S1). To assess gene completeness more thoroughly, we used TOGA2, a tool that generates whole-genome alignments to a high-quality reference genome (human: hg38) to identify conserved one-to-one orthologs and assess gene intactness, absence, or disruption^6^. *A. leucurus* displayed the greatest gene intactness, containing fewer missing or disrupted genes than any other squirrel species (Figure 1C and Data S2). Leveraging sequence alignments of conserved Sciuridae orthologs, we conducted a phylogenomic analysis that recapitulated established non-genomic phylogenies, recovering *A. leucurus* as the sister group to all other hibernators (Figure 1B).

**Figure 1.**
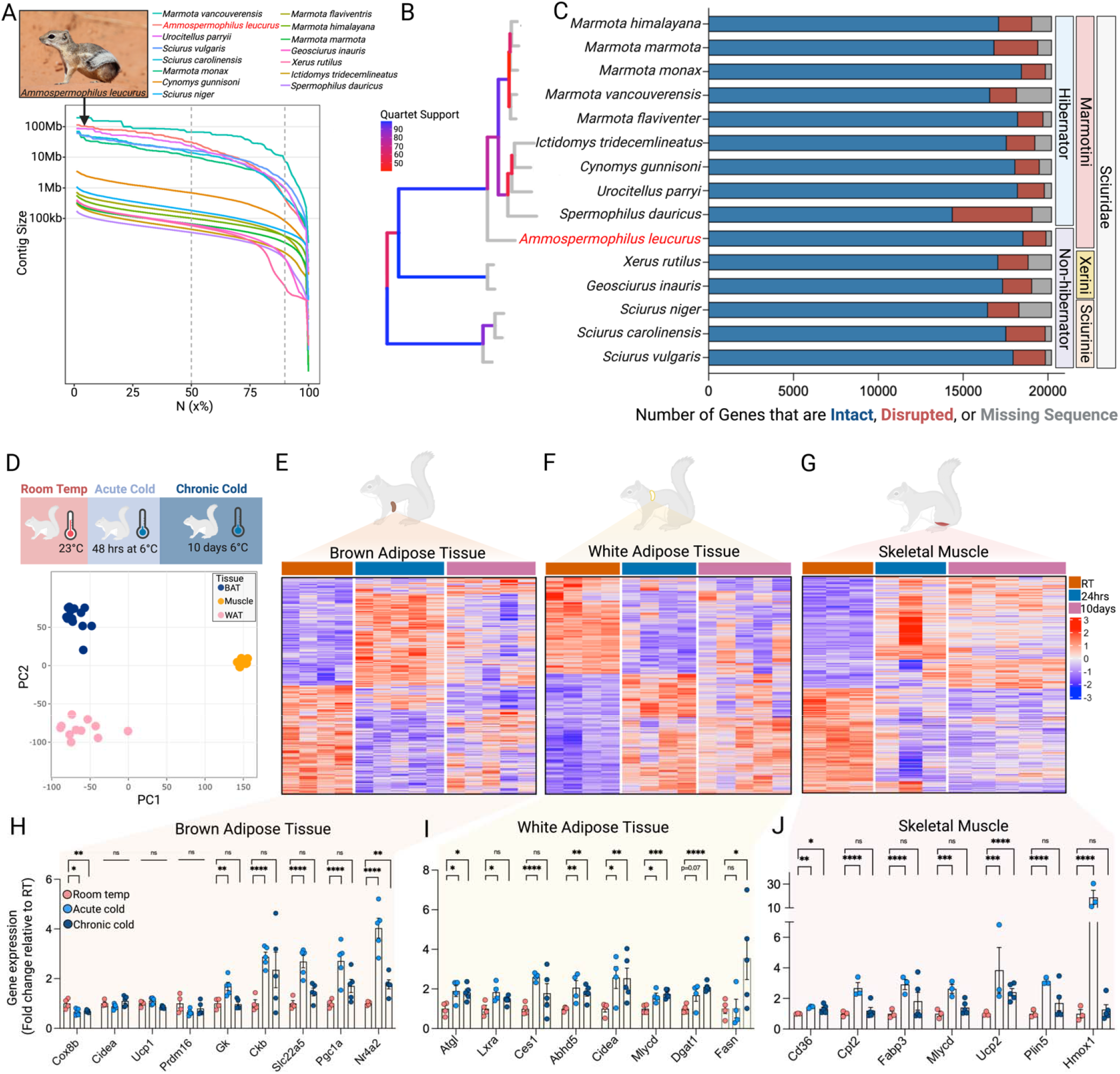
Comparative genomics and cold tolerance test in *A. leucurus*. **A**. Photo of a wild *Ammospermophilus leucurus* (top) and the contigN50 metrics across Sciuridae species (bottom). **B**. A phylogenetic tree of Sciuridae species included in the comparative genomics analysis. **C**. Corresponding TOGA gene intactness metrics for each species. Species are grouped into their genus and status as a hibernator or non-hibernator. **D**. Schematic of the cold exposure (top) and principal component analysis of all samples across treatments (bottom). A heatmap for all differentially expressed genes following cold exposure in brown adipose tissue (**E**.), white adipose tissue (**F**.), and skeletal muscle (**G**.). Genes with an adjusted *p* < 0.05 at either 24hrs or 10 days following cold exposure were included in the heatmap for each tissue. Genes of interest are shown as fold change relative to room temperature (RT) samples for brown adipose tissue (**H**.), white adipose tissue (**I**.), and skeletal muscle (**J**.). Adjusted *p*-values from DESeq2 are shown. ns = not significant, ^*^*p* < 0.05, ^**^*p* < 0.01, ^***^*p* < 0.001, ^****^*p* < 0.0001. N = 3-5 per group. The *A. leucurus* photo was taken by Renee Grayson near Las Vegas, Nevada, USA — CC BY 2.0, https://commons.wikimedia.org/w/index.php?curid=123681772.

To identify potential genomic drivers contributing to antelope ground squirrel thermal physiology, we scanned all orthologous genes across Sciuridae to identify predicted gene-inactivating mutations specific to *A. leucurus*. This comparative approach revealed ten gene losses, including *CLCA1* (chloride channel accessory 1), *TYMP* (thymidine phosphorylase), *MOGAT3* (monoacylglycerol O-acyltransferase 3), *PCDHB2* (protocadherin beta 2), *UROC1* (urocanate hydratase 1), *RTP5* (receptor transporter protein 5), *IGKV2D-29* (immunoglobulin kappa variable 2D-29), *CTAG1A* (cancer/testis antigen 1A*), OR10X1* (olfactory receptor 10X1), and *SMIM40* (small integral membrane protein 40) (Data S3). We confirmed these gene losses through sequence alignment with the corresponding mouse, cow, and elephant orthologs, and ruled out assembly base errors by inspecting HiFi read mappings. These gene losses are predicted to affect multiple tissues due to their expression patterns, including functions of the intestine (*CLCA1, MOGAT3)*, liver *(UROC1)*, brain *(RTP5)*, eye *(SMIM40)*, and testis *(CTAG1A*). Of these, loss of *UROC1* was notable because it encodes urocanase, which catabolizes urocanic acid. Given that urocanic acid is present on the skin^7^ and absorbs UV light^8^, UROC1 loss in this desert-dwelling species may reflect selection to increase or maintain urocanic acid levels.

Despite the identification of predicted gene losses with potential physiological relevance in *A. leucurus*, we did not find a specific loss that directly explains its altered thermogenic phenotype. This suggests that subtle regulatory changes, such as altered gene expression programs in classical thermogenic tissues, may more directly shape *A. leucurus* thermal physiology. To test this, we exposed laboratory-reared *A. leucurus* to either acute (24h) or chronic (10 day) cold (6°C) followed by bulk RNA sequencing of tissues that regulate shivering (skeletal muscle) and non-shivering thermogenesis (brown and white adipose tissue) (Figure 1D). Importantly, during chronic cold exposure, we gradually decreased ambient temperature to allow acclimation and minimize hypothermia risk (see Supplemental Material).

Brown adipose tissue (BAT), the main contributor to non-shivering thermogenesis^9^, demonstrated a rapid response following 24h cold, with 421 and 386 genes up-and down-regulated, respectively (Figure 1E and Data S4). While we observed increased gene expression of non-canonical thermogenic markers (futile creatine cycling: *Ckb*; futile lipid cycling: *Gk*) and mitochondrial regulators (*Pgc1a, Nr4a2, Slc22a5*), most canonical thermogenic markers showed no change (*Ucp1* and *Cidea)* or even decreased expression (*Cox8b)* (Figure 1H). This blunted thermogenic response persisted following 10 days of cold exposure, with thermogenic markers *Ucp1, Prdm16, Pgc1a*, and *Ckb* either remaining unchanged, returning to baseline, or decreasing below baseline levels (Figure 1H).

In marked contrast, subcutaneous white adipose tissue showed the strongest acute cold response, with 1,937 differentially expressed genes (DEGs) (Figure 1F and Data S4). These included increased expression of lipolytic genes *Ces1, Atgl, Lxra, Abhd5*, and *Mlycd* (Figure 1I). Although the number of DEGs dropped by nearly half after 10 days of cold exposure, white adipose tissue maintained robust transcriptomic remodeling, shifting toward a lipogenic profile (e.g., increased *Dgat1 and Fasn*) (Figure 1I). Notably, despite increased expression of some thermogenic markers (e.g., *Cidea*), we did not detect *Ucp1* expression in this depot (TPMs <1). Moreover, we observed no induction of genes previously linked to *Ucp1*-independent thermogenesis, including creatine cycling (*Ckb*/*Alpl*), calcium cycling (*Atp2a2*), or lipid cycling (*Gk*). These data suggest that the pronounced cold-induced remodeling of white adipose tissue does not contribute to direct heat production. Thus, the two adipose depots considered to produce heat via non-shivering thermogenesis display either a blunted response (brown fat) or limited thermogenic potential (white fat) at the level of gene expression.

These data suggest shivering (muscular) thermogenesis may be the primary heat source in *A. leucurus*. Consistent with this, acute cold exposure increased expression of skeletal muscle genes involved in fatty-acid uptake and transport (*Cd36, Fabp3, Plin5, Cpt2, Ucp2*, and *Mlycd*), supporting fatty-acid-fueled shivering (Figure 1G and 1J). Moreover, the most up-regulated gene was *Hmox1*, a key sensor and regulator of cellular oxidative stress^10^. Notably, in contrast to brown and white adipose tissue, skeletal muscle transcriptional responses continued to increase after 10 days of cold. Despite this sustained response, gene set enrichment analysis (GSEA) of DEGs at 10 days showed strong negative enrichment of programs linked to extracellular matrix organization, blood-vessel morphogenesis, and locomotion, suggesting a limited capacity for the remodeling required to sustain prolonged shivering thermogenesis (Data S5). Taken together, these data indicate that *A. leucurus* compensates for blunted adipose thermogenesis by relying primarily on muscular shivering. However, skeletal muscle shows a limited capacity for the remodeling needed to sustain prolonged shivering thermogenesis, such as enhanced vascularization and extracellular matrix remodeling. This constrained adaptive response may explain impaired cold tolerance of *A. leucurus* compared to its hibernating marmotine relatives. More broadly, our results highlight how evolutionary shifts in thermogenic programs may explain how closely related mammals thrive or fail under distinct thermal demands.

## Materials and Methods

### Genome sequencing

A spleen from a male *Ammospermophilus leucurus* specimen was used for genome sequencing. The sample, provided by the University of Washington Burke Museum (UWBM) Genetic Resources Collection, was UWBM # Mamm74590; Preparator J.R. Whorley #JRW 201. Detailed methods are described in the *SI Appendix*.

## Acknowledgements

We thank the University of Washington Burke Museum (UWBM) Genetic Resources Collection (Sharon Birks) for providing tissue samples used for genome sequencing. We thank Connie Zhao and Bin Zhang from the Rockefeller University Genomics Resource Center and Wei Wang of the Bioinformatics Resource Center for RNA-sequencing and processing. We thank UC Davis DNA Technologies Core for genomic sequencing and assembly. Figures were created in BioRender (https://BioRender.com).

## Author Contributions

LO and PC conceptualized the study; LO, MA, DB, and TC analyzed data; LO, MA, MH, RR, and GK designed experiments; LO, GK, and RR performed animal experiments; LO and PC wrote the manuscript and all authors reviewed and approved the final version.

## Data availability

Original RNA-sequencing data have been deposited in the Gene Expression Omnibus (GEO ID: GSE311886). The *A. leucurus* genome will be deposited upon publication.

