## Supplemental material for "Impaired non-shivering thermogenesis in the desert-dwelling antelope ground squirrel"

**This PDF file includes:**
Supporting text
Legends for Datasets S1 to S5
SI References

**Other supporting materials for this manuscript include the following:**
Datasets S1 to S5

### Extended Methods

#### Genome sequencing

*HMW DNA Extraction Method (HMW gDNA224)*
High molecular weight (HMW) genomic DNA (gDNA) was extracted from approximately 50 mg of spleen tissue using the Nanobind Tissue Big DNA kit, following the manufacturer's instructions (Pacific BioSciences, Menlo Park, CA). DNA purity was assessed by measuring absorbance ratios (260/280 = 1.84; 260/230 = 2.4) on a NanoDrop ND-1000 spectrophotometer. The final DNA yield (78.5 µg) was quantified using a Quantus Fluorometer with the QuantiFluor ONE dsDNA Dye assay (Promega, Madison, WI). The size distribution of the HMW DNA was analyzed using the Femto Pulse system (Agilent, Santa Clara, CA), which showed that 89% of the fragments were 10 kb or larger.

*Library preparation*
The HiFi SMRTbell library was constructed using the SMRTbell prep kit 3.0 (Pacific Biosciences, Menlo Park, CA; Cat. #102-182-700) according to the manufacturer's instructions. HMW gDNA was sheared to a target DNA size distribution between 18-20 kb using Diagenode's Megaruptor 3 system (Diagenode, Belgium; Cat. #B06010003). The sheared gDNA was concentrated using 1X SMRTbell cleanup beads provided in the SMRTbell prep kit 3.0 for the repair and a-tailing incubation at 37°C for 30 minutes and 65°C for 5 minutes, followed by ligation of overhang adapters at 20°C for 30 minutes, cleanup using 1X SMRTbell cleanup beads, and nuclease treatment at 37°C for 15 minutes. The SMRTbell library was subjected to a final bead cleanup using 1X AMPure PB beads (Pacific Biosciences, Menlo Park, CA; Cat. #100-265-900) in preparation for gel size selection.

*Gel size selection and HiFi sequencing*
Size selection was performed on the LightBench CF system for the removal of fragments 9 kb and smaller (Yourgene Health, Manchester, UK). The resulting library, with an average insert size of ~26.6 kb, was sequenced at the UC Davis DNA Technologies Core (Davis, CA) using one 25 M SMRT cell (Pacific Biosciences, Menlo Park, CA; Cat. #102-202-200) on the Revio system with SPRQ sequencing chemistry and a 30-hour movie time.

#### Genome assembly and structural annotation
PacBio HiFi data were assembled using Hifiasm[1] (v0.25.0-r726) with default parameters for HiFi assembly. Benchmarking sets of Universal Single-Copy Orthologs analysis[2] (v5.4.7) on the primary assembly showed 94.1% (S: 92.5%, D:1.6%) completeness using eukaryote_odb10 lineage and 91.8% (S:90.7%, D:1.1%) completeness using glires_odb10 lineage. Low-complexity regions, simple repeats, and interspersed repeats were masked in the primary assembly using the Dfam[3] database and the "-species sciurus" parameter in RepeatMasker[4] (v4.1.7-p1). Small RNA genes were left unmasked using the "-norna" option. The repeat-masked primary assembly was then used to annotate gene structures in Braker[5–11] (v3.0.3) with evidence from species-specific RNA-seq data, as well as mouse and marmot protein sequences.

#### Genome Annotation
Gene models generated by BRAKER3 were read into R, translated into protein sequences using the BioStrings package, and protein per transcript sequences exported as FASTA format for use in downstream tools. For functional annotation, eggNOG was implemented using default parameters with our protein FASTA file to provide predictions for functional annotations of genes including their putative GO, KEGG, and PFAM categories. For annotation of orthologous genes, reciprocal BLAST was run against mouse mm10 proteins retrieved from Ensembl with an E-value

cutoff of 1e-5, and the mutual best hit between gene models was selected. These results were combined with putative eggNOG orthologs to produce a final set of orthologous genes.

##### Genome alignment and gene annotation with TOGA

We used TOGA2 (Tool to infer Orthologs from Genome Alignments 2) (https://github.com/hillerlab/TOGA2), the next-generation version of the TOGA method[12] to assess gene completeness, provide coding gene annotations, and infer orthologs. Because TOGA relies on a whole-genome alignment, we first computed pairwise genome-alignment chains between human (hg38 assembly) as the reference species and *Ammospermophilus leucurus* as the query species, using lastz (parameters K = 2400, L = 3000, Y = 9400, H = 2000, default scoring matrix), axtChain (default parameters except linearGap = loose), RepeatFiller, and chainCleaner (default parameters except minBrokenChainScore = 75,000 and -doPairs)[13-16]. We then used TOGA2 with a new input annotation that combined the human GENCODE V46 annotation[17], the NCBI RefSeq All (curated and predicted) annotation release GCF_000001405.40-RS_2024_08[18] (both downloaded from https://hgdownload.soe.ucsc.edu/goldenPath/hg38/database/ on Oct 14, 2024), and MANE v1.4[19]. TOGA then infers orthologous gene loci using machine learning and alignments of intronic and intergenic loci, and annotates and classifies orthologous genes. To compare assembly completeness and base accuracy, we considered 18,430 genes that already existed in the placental mammal ancestor[20] and used the human-referenced TOGA classification to count how many genes have an intact reading frame, inactivating mutations, or missing sequence due to assembly gaps or assembly fragmentation.

##### Species Tree Inference

For a rigorous and reproducible species tree inference, all one-to-one orthologs annotated by TOGA2 with at least one intact or fully intact transcript were selected. For each gene, a single transcript was chosen based on overall intactness across the selected assemblies, resulting in 5,476 transcripts. These transcripts were aligned across species using PRANK v.250331[21], with the human genome (hg38) as the reference and the species tree from[22] as the guiding phylogeny. Gene trees were inferred for each transcript alignment using IQ-TREE 2 v2.4.0[23] with automatic model selection. The individual gene trees were then concatenated and reconciled into a single species tree using ASTRAL-IV v1.23.4.6, implemented in the ASTER[24] package. Branch supports were calculated based on the frequency of each winning quartet across the individual gene trees, considering the three possible quartet topologies.

##### Gene Loss Identification

TOGA2 identifies gene losses by examining the presence and pattern of inactivating mutations and classifies each gene into one of the following categories: Intact (I), Partially Intact (PI), Uncertain Loss (UL), Missing (M), Lost (L), or Paralogous (PG). Gene loss status was retrieved for all target species and filtered to retain only those genes categorized as L in the focal species (*Ammospermophilus leucurus*), while being FI, I, PI (the statuses indicating the absence of inactivating mutations) in the other species. The presence of other categories (e.g., UL) was tolerated in up to two background species. All candidate genes were then manually curated by reviewing the inactivating mutations to identify false uncertain losses that were more likely to represent intact genes. The final candidate genes were further validated by reassessing their loss status with TOGA2 run against three additional reference genomes: mouse (*Mus musculus*), cow (*Bos taurus*), and elephant (*Elephas maximus indicus*).

To account for potential base errors in our assembly, which can mimic false gene losses[25,26], HiFi *Ammospermophilus leucurus* sequencing reads were mapped to the genome assembly with minimap2[27]. We then manually inspected the lost genes to verify even read coverage across the gene locus and to ensure that the observed inactivating mutations are supported by the raw HiFi reads. For all gene losses reported in this manuscript, we identified at least one homozygous mutation in the sequenced individual, for which all or nearly all aligned reads support the genome assembly.

**Cold exposure**

Animal studies were approved by the ethics committee of the University of New Orleans (approval number: IACUC 25-001). A group of eighteen male and female adult *Ammospermophilus leucurus* were subdivided into three groups of six. In each group, squirrels were individually housed in polypropylene cages (36 cm length, 24 cm width, 19 cm height) lined with wood shavings. Purina rodent chow (laboratory rodent diet 5001) and drinking water were provided *ad libitum*. Squirrels were provided with three to five cotton balls throughout the experimental treatment as nesting material. The control group remained at room temperature (23°C) throughout the study. The second group was exposed to cold (6°C) for 24 hours, and the third group was exposed to cold for 10 days. For the 10-day group, a gradual decline in ambient temperature was used, decreasing from 23°C to 11°C over the first two days and then held constant at 6°C from day three until day 10. Cold exposure was produced in a ventilated refrigerator with a thermometer placed inside for continuous temperature monitoring. All animals were inspected daily, with no signs of abnormal behavior that would warrant termination of the study. Following the respective treatments, squirrels were euthanized by $CO_2$ inhalation and tissues were collected. White adipose samples were collected from the interscapular adipose depot. While classically considered a brown fat depot, this depot takes on a white adipose phenotype across many squirrel species; this was validated by the absence of *Ucp1* gene expression. Brown fat was collected from the axillary depot, and skeletal muscle was collected from the hindlimb (gastrocnemius). All tissue samples were cleaned and rapidly frozen in liquid nitrogen.

**RNA extraction and sequencing**

Frozen tissue was homogenized in a bead beater with Trizol (ThermoFisher Scientific: 15596026). RNA was separated via the Trizol-chloroform method. The subsequent RNA fraction was combined with 70% ethanol and processed with the Qiagen RNeasy Plus Micro Kit (Qiagen: 74034), including DNase incubation treatment. A total of 100 ng of RNA was used to generate RNA-Seq libraries using the Illumina stranded mRNA kit (Catalog #: 20040534), following the manufacturer's protocol. Libraries prepared with unique dual indexes were pooled at equal molar ratios. The pool was sequenced on an Illumina NextSeq 2000 P3 flowcell using NextSeq Control Software v1.7.1.46395 to generate 100 bp (or 50 bp) paired-end reads, following the manufacturer's protocol (Document #: 200027171 v04).

**Data analysis for RNA-sequencing**

Transcript abundance was determined from FASTQ files using Salmon (v1.8.0) and Ensembl reference transcript sequences[28]. Transcripts per million (TPM) values were calculated using the tximport R Bioconductor package (v1.22.0)[29]. Differential gene expression analysis was performed with the DESeq2 R Bioconductor package (v 1.44.0)[30]. For paired samples, differences in gene expression across time points or conditions were calculated while controlling for each subject in the linear model. Gene counts were transformed with the vst function from DESeq2 to apply a variance-stabilizing transformation, and z-scores of these normalized values for the indicated gene sets were visualized with heatmaps generated using the ComplexHeatmap R package (v1.0.12)[31]. For GSEA, the Wald statistic from the DESeq2 analysis was used to generate the ranked list of genes. The clusterProfiler R Bioconductor package (v4.12.6) was then used to perform GSEA to determine the enrichment of the Gene Ontology Biological Process gene sets sourced using the msigdbr R package (v25.1.1)[32-34].

#### Funding

LO was supported through the Simons Foundation Postdoctoral Fellowship and a BFTDF pilot award. PC was supported by the NIDDK (RC2 DK129961) and Leducq Foundation.

#### Declaration of AI-assisted technologies in the writing process

This manuscript benefited from limited use of ChatGPT (OpenAI) to assist with initial grammatical editing and improving conciseness; all scientific content and interpretations were generated and verified by the authors.

**Data S1 (separate file). *A. leucurus* genome assembly statistics.**

**Data S2 (separate file). Squirrel species and genome assemblies used for comparative analysis.**

**Data S3 (separate file). Gene loss identified within *A. leucurus*.**

**Data S4 (separate file). Identification of differentially expressed genes following cold exposure in *A. leucurus*.**

**Data S5 (separate file). Gene set enrichment analysis for biological process of skeletal muscle.**
